# Photodynamic Therapy for Deep Organ Cancer by Implantable Wireless OLEDs

**DOI:** 10.1101/2024.11.18.624193

**Authors:** Yujiro Itazaki, Kei Sakanoue, Katsuhiko Fujita, Izumi Kirino, Kazuhiro Eguchi, Yutaka Miyazono, Ryoichi Yamaguchi, Takasumi Tsunenari, Takao Sugihara, Kenji Kuwada, Naoki Kobayashi, Tsuyoshi Goya, Katsuyuki Morii, Hironori Tsujimoto, Yuji Morimoto

**Affiliations:** Department of Surgery, National Defense Medical College, Saitama, Japan; Pleiades Technologies LLC., Fukuoka, Japan; Institute for Materials Chemistry and Engineering, Kyushu University, Fukuoka, Japan; Division of Hepato-Biliary-Pancreatic Surgery and Transplantation, Department of Surgery, Graduate School of Medicine, Kyoto University, Kyoto, Japan; Kumamoto Industrial Research Institute, Kumamoto, Japan; NIPPON SHOKUBAI CO., LTD., Osaka, Japan; Nippon Shokubai Research Alliance Laboratories Osaka University, Osaka, Japan; Department of Physiology, National Defense Medical College, Saitama, Japan

**Keywords:** Metronomic photodynamic therapy, Organic light-emitting diode (OLED), Wirelessly powered, Deep organ cancer therapy, Apoptosis

## Abstract

Metronomic photodynamic therapy (mPDT) is a method of continuously delivering low-intensity light to a cancer lesion. Since high light intensity is not required, the light device can be miniaturized and implanted in the body. Nevertheless, for clinical implementation, it is essential to develop a light source apparatus that can effectively illuminate the entire tumor and operate stably without damaging internal organs. To address this issue, an organic light-emitting diode (OLED) optical device that has an area large enough to cover the entire lesion and is powered by a wireless power supply has been developed. The device is thin and lightweight and it functioned stably in the abdominal cavity of rats for more than two weeks without causing foreign body reactions. By covering the tumor site of rat liver cancer with this device and continuously illuminating the tumor with appropriate administration of a photosensitizer, the entire tumor under the emitting surface was eliminated through apoptosis without leaving any residual tumor margins. This is the first report of a robust antitumor effect on parenchymal organ cancers by mPDT using a biocompatible, wirelessly powered OLED device, which represents a potential new deep organ cancer therapy.

## 1. Introduction

Photodynamic therapy (PDT) uses photosensitizer (PS) activation by intense, brief light exposure (>100 mW/cm^2^ for ∼10 min) to generate reactive oxygen species, resulting in death of cancer cells[1]. PDT is clinically approved for treatment of superficial cancers in organs such as the esophagus, lung, stomach, cervix, and brain. Tumors can be irradiated directly by a light source or through an optical fiber.

Metronomic PDT (mPDT) is a newly developed method using repeated PS administration with extended low-intensity light[2]. Unlike conventional PDT with light intensity of more than 100 mW/cm^2^, mPDT uses less than 1 mW/cm^2^ of light intensity. This reduces thermal damage risks and allows the light source to be miniaturized and implanted in the body[3]. Besides, extended illuminating time in mPDT can continuously inhibit tumor growth. The authors previously developed a wirelessly powered LED device for mPDT and proved its efficacy against subcutaneous tumors[4] [5].

Despite promising developments, mPDT using an implantable device for tumors in deep organs, such as the liver and pancreas, has not yet been realized. This is because no device has met the requirements of 1) being able to deliver light to the entire tumor area, 2) being powered wirelessly from outside the body to maintain constant light intensity, 3) being driven within the body for a sufficient duration for treatment, and 4) not causing immunological problems or heat damage. To address these challenges, the authors have developed a wireless light-emitting device using an OLED that emits light homogeneously over a large surface, covering the entire tumor, and applied it for mPDT in deep organs.

The newly developed OLED device is compact and thin enough to be implanted anywhere in the body. It can deliver light to the entire lesion in a deep organ, overcoming challenges of conventional light delivery. A stable wireless power supply, achieved through a tuned system, ensures consistent light emission, highlighting the successful application of wireless power transmission techniques. A parylene coating further enhances the device’s protection and biocompatibility of the device.

Experiments using a rat model of orthotopic hepatoma showed that the newly developed mPDT system has remarkable efficacy in the treatment of deep organ cancer and has the potential to overcome the limitations imposed by conventional PDT. This study is the first study of a robust anti-tumor effect on parenchymal organ cancer by mPDT using a biocompatible, wireless-powered OLED device, and it represents a potential step toward clinical application.

## 2. Results

### 2.1 A Wirelessly Powered OLED Device

A conceptual diagram of mPDT using the new device is shown in Figure 1. The implantable wirelessly driven OLED device is placed on the tumor surface of the deep organ. The tumor is continuously illuminated by the OLED with repeated administration of the PS.

**Figure 1.**
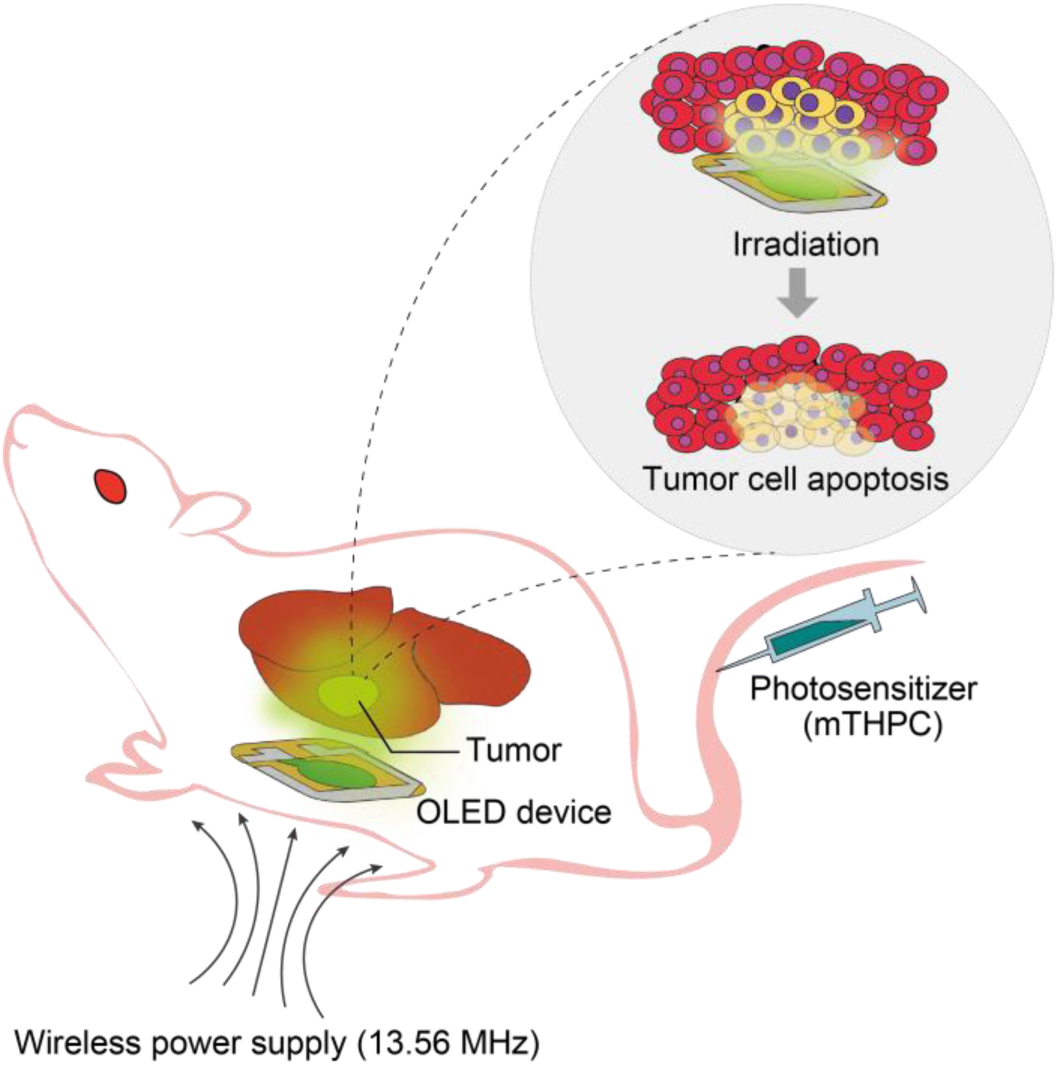
Concept of mPDT using the OLED device. After the OLED device has been attached to the animal tumor, the OLED emits light by wirelessly power while the PS is repeatedly administered to the animal, inducing cell death in the tumor.

The structure and various properties of the implantable wireless OLED device are as follows. The device measures 15.0 (width) × 16.0 (length) × 1.8 (thickness) mm and weighs 0.6 g (Figure 2a, b). The emitting area is circular with a diameter of 8 mm, covering the entire cancer lesion. The OLED element is composed of a transparent electrode, electron injection layer, electron transport layer, light emission layer, hole transport layer, hole injection layer, and aluminum[6] deposited on a 50-micrometer polyethylene terephthalate substrate. To this OLED element, a power-receiving coil, matching circuit unit, and ferrite sheet are added to form a wirelessly powered OLED device (Figure 2c). The power-receiving coil (13.6 mm φ) is tuned to the resonant frequency at 13.56 MHz and is electromagnetically shielded with a ferrite sheet to protect against a reduction in the Q-factor due to interference (Supplementary Table 1). The Q-factor reflects how sharply energy is stored in the resonant circuit, and it also indicates the effectiveness of energy transmission.

**Figure 2.**
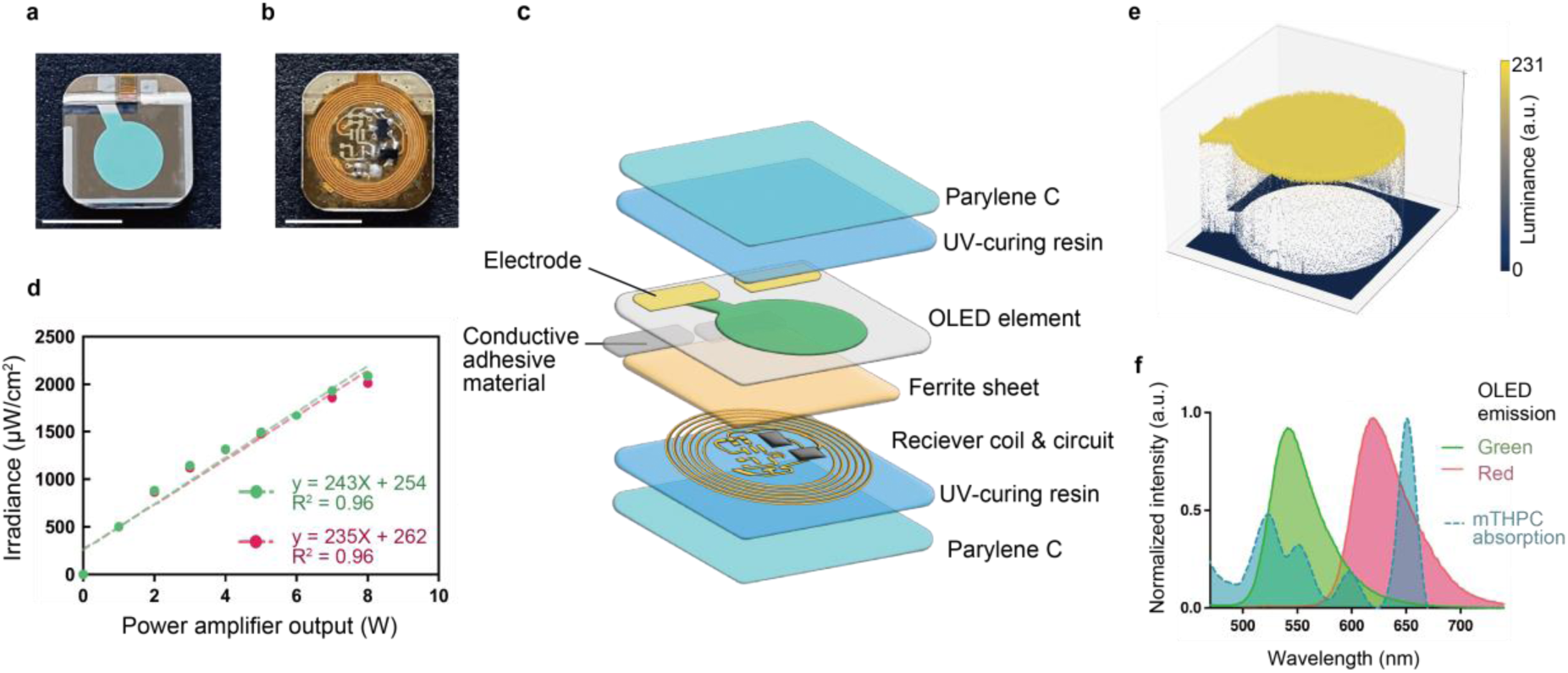
Implantable OLED device with its structure and properties. Both sides of the OLED device: the OLED element is housed on one side (a) and the power-receiving coil (2 layers) is on the opposite side (b). Scale bar = 10 mm. c) Stratified structures of the OLED device. d) Irradiance – operating power characteristics of the OLED devices (green, red). e) Luminance mapping at the OLED emitting surface. f) Absorption spectrum of the PS (mTHPC; 5,10,15,20-tetrakis (3-hydroxyphenyl) chlorin) and emission spectra of the OLED devices (green, red). a.u, arbitrary units.

To ensure waterproofing and biocompatibility, the OLED device is coated with UV-curing resin and Parylene C[7] (Figure 2c). The parylene coating, which is made through chemical vapour deposition and is known for its low pinhole content, serves as an insulator for organic transistors and as an encapsulation layer for flexible devices[8].

The use of an OLED has led to increased uniformity in luminescence across the surface (variation of luminance over the entire luminous area: ± 0.8%) (Figure 2e, Figure S1b) as well as high linearity of luminous intensity with respect to the operating power (correlation coefficient > 0.979) (Figure 2d), with green and red OLEDs having virtually identical characteristics.

As a PS for efficient photoexcitation in the emission spectrum of the OLED, temoporfin (5,10,15,20-tetrakis (3-hydroxyphenyl) chlorin; mTHPC)[9] was used. As shown in Figure 2f, the absorption spectrum of mTHPC overlaps with the emission spectrum of the green OLED (∼540 nm) and red OLED (∼620 nm), indicating that of the mTHPC is efficiently excited by the respective OLEDs.

### 2.2 Stably Working OLED Device *In vivo*

Metronomic PDT requires continuous illumination with low-intensity light over a long period of time. Thus, the implanted light source device must emit light persistently and stably, even when the animal is free to move around.

A novel power transmission platform was developed to provide a constant power supply to ensure that the OLED device remains activated when the animals are moving freely in the cage (Figure 3a, b). The transmitter was designed to be large enough for placing commercially available rodent cages (bottom dimensions: 206 mm (W) × 365 mm (L)) by adopting a uniaxial Helmholtz coil antenna that ensures uniform power distribution in the vertical direction. As a result, the irradiance of the OLED remained within a narrow range of 565 ± 100 µW/cm^2^ (mean ± SD) when the vertical height level of the OLED position was varied between 35 mm and 135 mm (Figure 3c,d,e). These data show that the OLED placed on the liver of a rat can be driven stably with a variability of ±18%, no matter what position or posture the rat in the cage is in.

**Figure 3.**
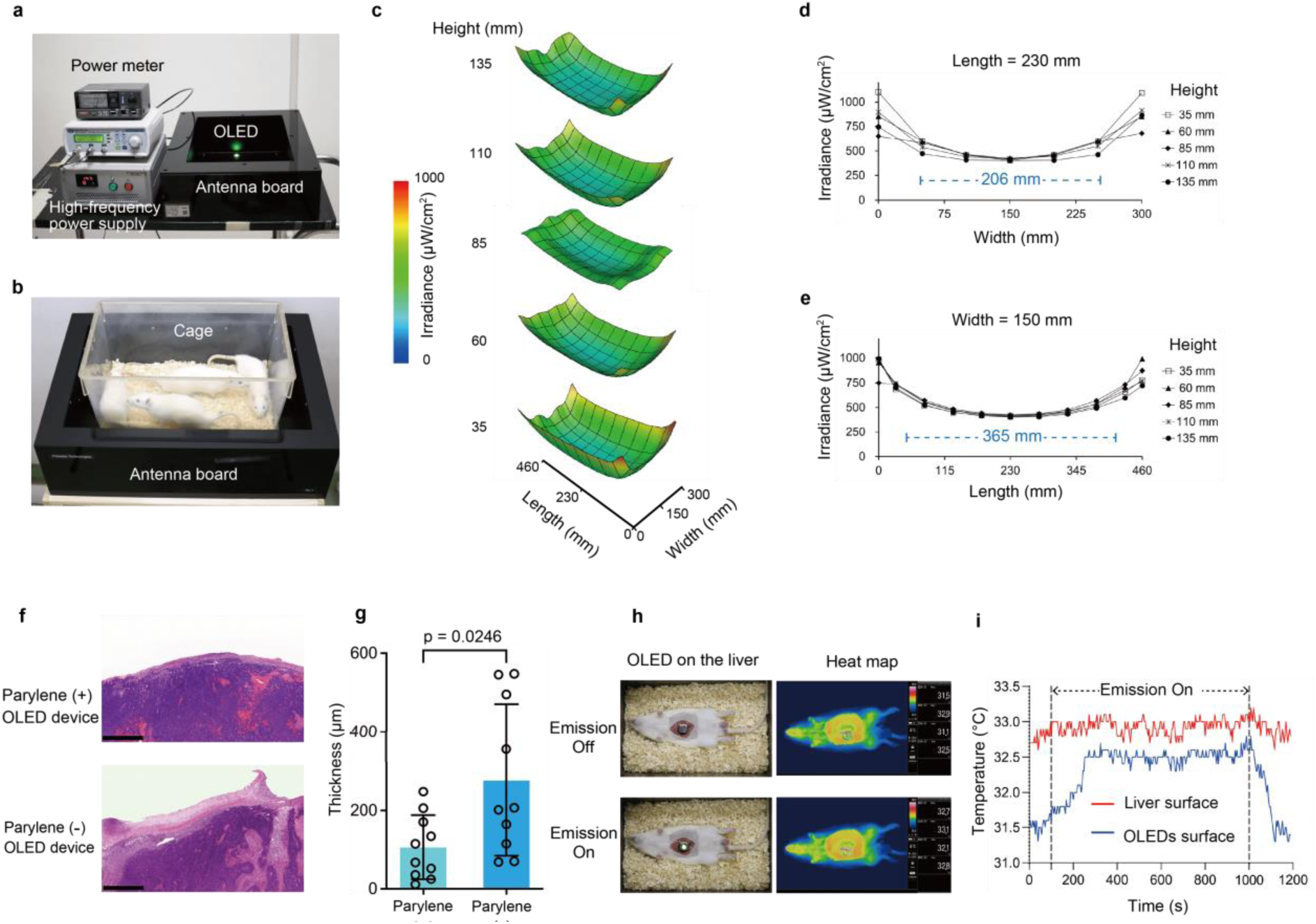
Suitability of the OLED device for *in vivo* applications. a) Power transmission platform system. b) Animals moving around in a cage on the antenna board. c) Three-dimensional distribution of irradiance from the green OLED when driven by the power transmission platform with the power setting of 8 W. The measured height range between 35 mm and 135 mm corresponds to the height of the OLED attached to the liver of a rat lying on the cage floor (35 mm) to the height of the OLED when the rat stands up (135 mm). Irradiance distributions of the OLED as seen in the cross section cut at 230 mm in the longitudinal direction of Figure 3c (d) and that at 150 mm in the width direction of Figure 3c (e). The blue dashed lines indicate the width (206 mm) (d) and length (365 mm) (e) of the cage bottom. Sections of the tumor in the liver 1 week after attachment of the green OLED device (f) (H&E staining, scale bar = 1 mm) and thickness of the fibrous tissue formed (g) (n = 10 in each). Fibrous thickening can be seen on the surface of the tumor in the liver with the parylene (-)-OLED device, while it is slight in the liver with the parylene (+)-OLED device. Heat map (h) and temperature change (i) when the OLED affixed to the liver surface was working. The temperature rise of the OLED device is at most 1°C, and no temperature rise is seen in the liver near the OLED device. A green OLED was used and the transmitter power setting was 8 W.

When the implantable device is used in the body, water sealing is required for continuous light emission. Water sealing by Parylene C and resin allowed the OLED device to be operated in saline solution. The degradation of light intensity was less than 2.7% even after 35 days of continuous light emission at 8 W of transmitter power (Figure S2a,b,c). Furthermore, OLED devices implanted in the rat abdominal cavities were confirmed to work for more than 2 weeks.

Although the implantation of an electrical device in the body causes foreign body reactions[10], a parylene coating can reduce unfavorable reactions[11] and was thus used in this OLED device. Two groups were studied to evaluate the reactions: one group with only UV-curing resin coating and another group with both resin and Parylene C coatings. One week after implantation of the devices on tumors in the livers, capsule thickness was analysed. Parylene C-coated devices showed a significantly lower foreign body reaction with a much smaller capsule thickness than that of the resin-only-coated devices (91.7 μm vs. 204.5 μm, n=10 respectively) (Figure 3f, g).

When electronic devices are used *in vivo*, the heat generated during operation can cause thermal damage. However, heat generation from the working OLED device was negligible (approximately 1.0°C) (Figure 3h,i), and no thermal damage to the liver tissue where the device was placed was seen.

### 2.3 Metronomic PDT Using the Implantable OLED Device

Having confirmed that the wirelessly powered OLED device can be used safely *in vivo*, its anti-tumor effect was then evaluated in an orthotopic hepatoma model.

An orthotopic rat hepatoma model was established by inoculating N1S1 cancer cells into subserous tissue of the left hepatic lobe. Four days after the inoculation, a PS (mTHPC) was administered intravenously (0.5 mg/kg). At that time point, the tumor size was 20 - 30 mm^3^. The OLED device was attached to the liver with adhesive drops at the corners of the device (4 points) so that the emitting surface covered the tumor (Figure S3a).

The OLED device attached to the liver emitted light via a wireless power supply (Supplemental Movie 1) and continued to emit light stably under conditions of body motion (Supplemental Movie 2) after abdominal closure.

To estimate the total amount of light energy received by the tumors exposed to the OLEDs during mPDT, movement of the animals in a cage within the antenna board was traced. By calculating the time the animals spent in each area and the light emitted in each area, the light intensity per second from the OLED was found to range from approximately 493 to 643 μW/cm^2^ when the transmitter power setting was 8 W. Thus, the cumulative light energy over a period of 48 hours was estimated to be approximately 85-111 J/cm^2^ (Figure S4). The characteristics described above were considered to show similar results for the green and red OLEDs, because the irradiance relative to the operating power was virtually identical (Figure 2d).

### 2.4 Eradication of the Whole Tumor

An investigation was conducted to prove the potent anti-tumor effect of mPDT using an OLED device. First, enhancement of the therapeutic effect by repeated administration of the PS was investigated and then the superiority of different illumination wavelengths on the therapeutic effect was examined.

#### 2.4.1 Repeated PS Administration Enhances Therapy

To demonstrate the superiority of repeated PS administration in the therapeutic effect in mPDT, three experimental groups were studied using green OLEDs according to the protocol shown in Figure 4a: (i) a control group implanted with a non-luminescent dummy device and receiving PS administration twice, (ii) an mPDT group implanted with a luminescent device and receiving PS administration once, and (iii) an mPDT group implanted with a luminescent device and receiving PS administration twice.

**Figure 4.**
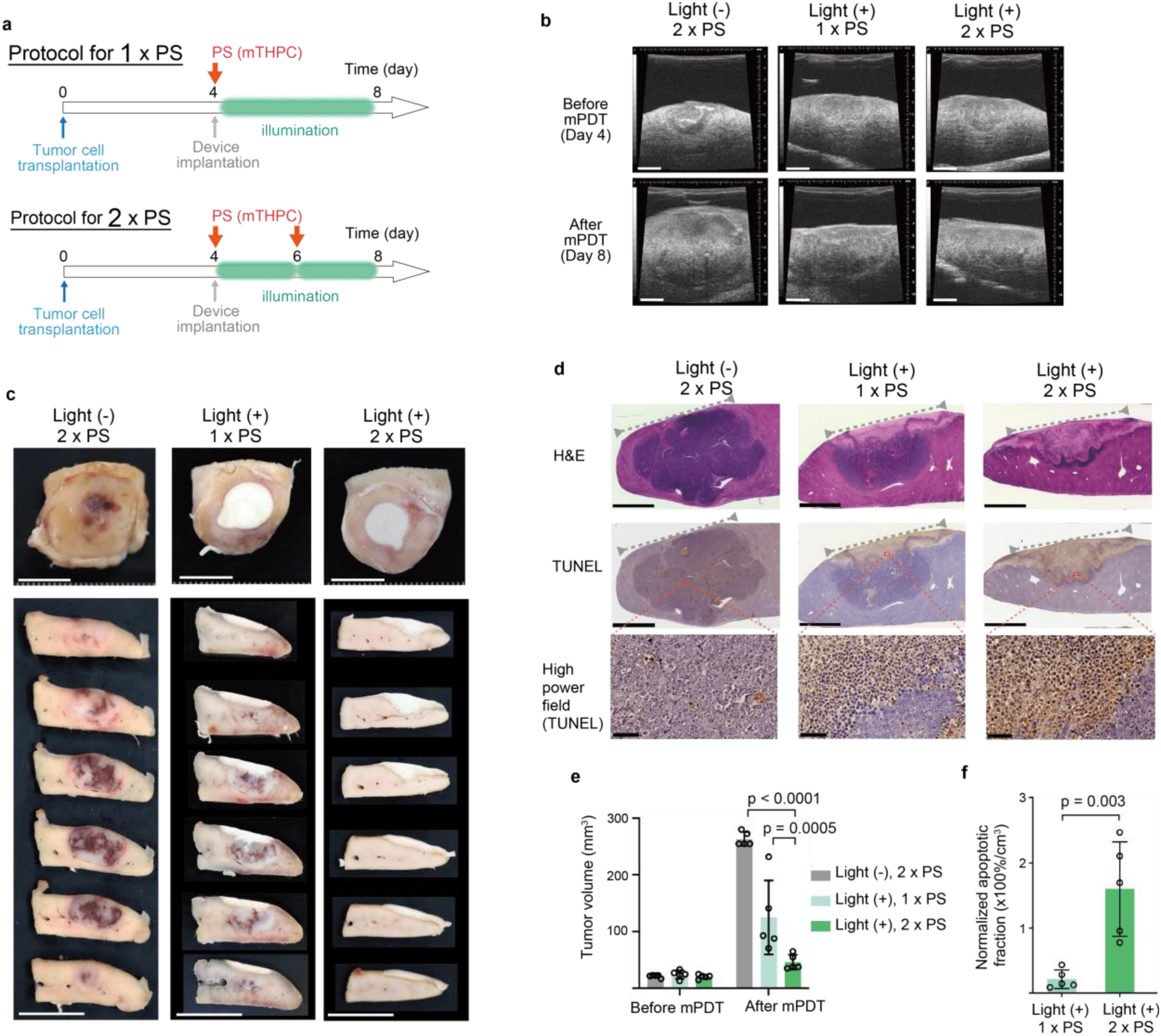
Anti-tumor effects of mPDT using the OLED device and enhancement of the effects by repeated PS administration. a) Experimental protocol for mPDT with change in the frequency of PS administration. Rats were inoculated with cancer cells in the liver lobe and, following an interval of four days, the OLED device was implanted. The PS was then administered via the caudal vein, and the tumor was illuminated for 2 days. In the protocol of one-time use of the PS, an additional 2 days of illumination was executed without additional administration of the PS, whereas in the protocol with two-time use of PSs, an additional 2 days of illumination was executed with additional administration of the PS. b) Ultrasound images before and after mPDT. The echo value of the tumor tissue is slightly lower than that of the surrounding liver tissue, which allows detection of the tumor border. c) Liver surface after removal of the OLED after mPDT (top) and crosssection of the liver specimen sliced at 2-mm intervals in the ventrodorsal direction (bottom). The mPDT-treated liver surface shows distinct pallor changes. d) Cross-sectional views of tumors after mPDT: H&E staining (top), TUNEL staining (middle), and high magnification image of the TUNEL staining (bottom). *Dashed lines with arrowheads* on the tumor surface indicate the position of the OLED emitting surface. e) Contrastive analysis of tumor volume changes in each experimental group before and after mPDT (n=5, each). f) Normalized apoptotic fractions, reflecting cell death relative to tumor volume (n=5, each). Scale bars: 2 mm in (b), 10 mm in (c), 0.05 mm in high magnification and 2.5 mm in other histopathology images for (d). The transmitter power was set at 8 W.

Tumor volume was assessed using ultrasound imaging based on the dimensions of the tumor before and after treatment (Figure 4b). The rats with the luminescent device (groups (ii) and (iii)) had a significantly smaller tumor volume than that in the control rats (Figure 4c,d,e). Furthermore, administration of the PS twice (group (iii)) induced significantly higher anti-tumor effects than did administration of the PS once (group (ii)), with mean volumes at the end of treatment being 263 mm^3^ in group (i), 106 mm^3^ in group (ii), and 37.7 mm^3^ in group (iii) (Figure 4e), emphasizing the benefits of additional administration of the PS. In the mPDT groups (groups (ii) and (iii)), whitening of tissues in contact with the OLED luminescent surface was seen and cross-sectional observation showed that the entire cancer lesion was whitened (Figure 4c).

Histopathological images showed significant apoptotic changes in the mPDT-treated groups (Figure 4d): nuclear abnormalities (pyknosis, karyorrhexis) and an increase in TUNEL-positive cells at sites where an antitumor effect was observed. When compared to the mPDT with one-time administration of the PS, there was more extensive cell death in the two-time group, resulting in a higher apoptotic fraction (Figure 4f). In addition, the therapeutic effect was also steadily expanded to the tumor margins, with no residual tumor cells in the marginal area.

#### 2.4.2. Illumination Wavelengths and Treatment Efficacy

In order to determine the predominant OLED emission wavelength in tumor therapeutic efficacy, the following three experimental groups (all with the PS administered twice) were studied according to the protocol shown in Figure 5a: (i) a control group implanted with a non-luminescent dummy device, (ii) a group implanted with a green luminescent device and (iii) a group implanted with a red luminescent device.

**Figure 5.**
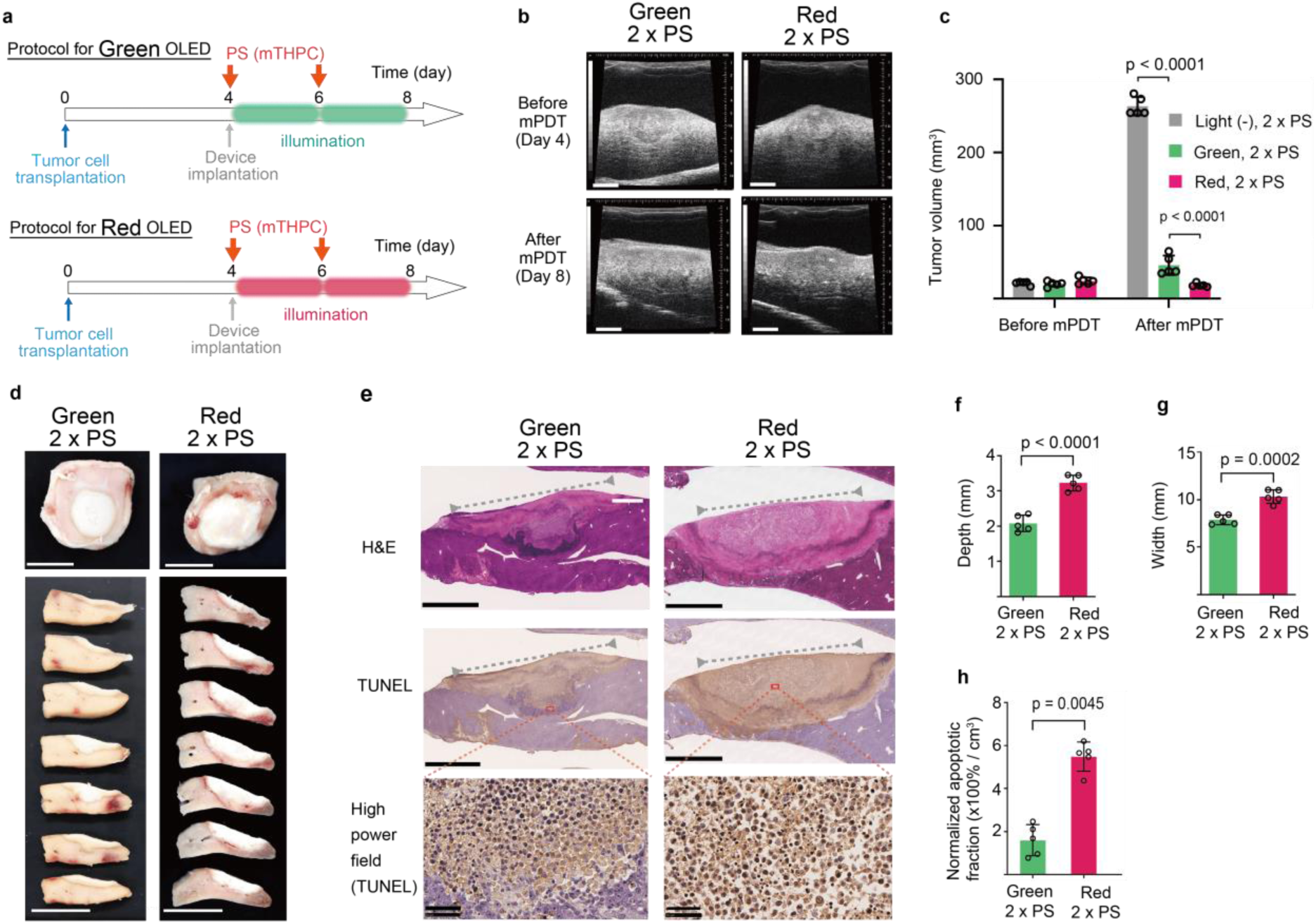
Augmentation of anti-tumor effects with red OLEDs. a) Experimental protocol for mPDT using OLED devices with different emission wavelengths: procedure the same as “Protocol 2 x PS” in Figure 4a with only the emission wavelength of the device to be implanted being different. b) Ultrasound images before and after mPDT. c) Tumor volume changes in each experimental group before and after mPDT (n=5, each). d) Liver surface after removal of the OLED after mPDT (top) and cross-section of the liver specimen sliced at 2-mm intervals in the ventrodorsal direction (bottom). The mPDT-treated liver surface shows distinct pallor changes. e) Cross-sectional views of tumors after mPDT: H&E staining (top), TUNEL staining (middle), and high magnification image of the TUNEL staining (bottom). *Dashed lines with arrowheads* on the tumor surface indicate the position of the OLED emitting surface. f) Deepest reach of the cell death zone in the vertical direction of the emission plane. g) Maximum reach of the cell death zone in the horizontal direction of the emitting plane. h) Normalized apoptotic fractions, reflecting cell death relative to tumor volume (n=5, each). Scale bars: 2 mm in (b), 10 mm in (d), 0.05 mm in high magnification and 2.5 mm in other histopathology images for (e), 2.5 mm in (i). The transmitter power was set at 8 W for both green and red OLEDs.

Metronomic PDT with red OLEDs showed a significantly larger therapeutic area than that with mPDT using green OLEDs (Figure 5d, e). Specifically, the red OLEDs significantly increased the therapeutic effect in both vertical (Figure 5f) and horizontal (Figure 5g) directions based on the emitting area, especially in the horizontal direction across the emitting area (Figure 5e, g). As a result, mPDT with the red OLED induced more tumor cell death (Figure 5e, h) and the tumor volume was significantly smaller than that when mPDT with the green OLED was used (Figure 5c).

To assess potential adverse effects, blood samples were collected before and after mPDT to measure hepatic enzymes. Aspartate aminotransferase (AST) and alanine aminotransferase (ALT) levels during treatment showed no significant difference between the two-time mPDT group and control group (Figure S5a, b). Similarly, changes in body weight over time during treatment were not significantly different between the groups (Figure S5c).

## 3. Discussion

To realize mPDT for deep organ cancer, a thin, lightweight, biocompatible, and wirelessly powered OLED device that can cover the tumor surface and illuminate the entire tumor has been newly developed. Using this device, mPDT successfully reduced tumor size in an orthotopic rat hepatoma model.

The use of the OLED device has made it possible to distribute light homogeneously and to cover the marginal boundaries of the tumor. This effectively activates localized PSs and produces massive anti-tumor effects. Since tumors grow by replacing adjacent healthy tissue[12], illuminating the tumor margin is crucial. Including the surrounding healthy tissue in the illumination helps reduce tumor regrowth[13], [14].

The OLED device developed in this study was coated with Parylene C not only for waterproofing[15] but also for biocompatibility[16], and it was shown to have sufficient immunological biosafety for remaining in the body long enough for deep cancer treatment. Since the rise in temperature during treatment was only 1°C at most, thermal damage to the normal tissue surrounding the tumor, which is a concern when using stronger light such as laser[17], was not observed.

The spread of cancer cells through surgical incisions and punctures of the tumors, one of the major concerns in cancer treatment[18], can be minimised because the OLED device is simply attached and fixed to the surface of the tumor. This device maintains a more stable position relative to the lesion than do conventional endoscopic methods, as it ensures consistent alignment between the light source and the tumor, resulting in a more stable and effective light delivery directly to the tumor. Therefore, therapeutic effects can be expected in long-term treatment.

Metronomic PDT with administration of the PS twice efficiently disintegrated tumor cells, leading to tumor tissue atrophy. This suggests that light to active tumor cells lying deep in the atrophic layer was increased and the therapeutic effect was expanded in a deeper direction. Therefore, mPDT with sophisticated control of PS administration can surmount the limitation in conventional PDT, which has a therapeutic effect only near the surface, and mPDT expands the therapeutic reach into deeper layers of the cancer tissue.

This study confirmed the superiority of the therapeutic effect of red light compared to green light. In particular, the therapeutic effect of mPDT using the red OLED was shown to be expanded in both deep and horizontal directions with respect to the emitting surface. Red light penetrates deeper into the tissue than does green light [19] and it allows a wider and more uniform distribution of light, including horizontally. Moreover, the red OLEDs used showed a stronger horizontal light distribution than did the green OLEDs (Figure S7), which may have especially enhanced the horizontal expansion. Although the superiority of red light is known from previous PDT studies, the homogeneous illumination and fixed positioning of the light source contributed to the enhancement and expansion of the anti-tumor effect of red light, highlighting the advantages of the surface-emitting OLEDs.

## 4. Conclusion

A novel device for treating deep organ cancer with mPDT has been developed. This thin, lightweight, and biocompatible OLED device is wirelessly powered and can illuminate entire tumors. Its effectiveness was proven in a rat hepatoma model by significantly reducing tumor size. This study is the first study to show that mPDT with a wirelessly powered, tumor-covering OLED device is effective for parenchymal organ cancers. The mPDT system could transform the treatment of deep cancers that are challenging to treat with conventional PDT. Future enhancements will include integration with an external, portable power unit to precisely control illumination timing, potentially revolutionizing cancer treatment and facilitating home-based care.

## 4. Experimental Section

### Fabrication of the Wirelessly Driven OLED Device

The layer configuration of an OLED element is as follows: transparent electrode / electron injection layer / electron transport layer / emission layer / hole transport layer / hole injection layer / aluminum[6]. Each layer was sequentially constructed on polyethylene terephthalate substrates using a vacuum process. The fabricated structure was then exposed to ambient air and sealed with a sealing film. To this OLED element, a power-receiving coil, matching circuit unit, and ferrite sheet were added to form a wirelessly powered OLED device (Figure 2c). The power-receiving coil was constructed by etching copper foil formed on both sides of a polyimide substrate into a spiral pattern. The diameter of the coil was set at 13.6 mm, and two layers of coils with a wiring width of 0.2 mm and number of turns of 7 were made to maximise resonance at 13.56 MHz most efficiently. The matching circuit and the use of ferrite sheets were tuned to optimise the Q-factor. After integration of the OLED element and receiver coil, they were coated with a UV-curable resin made of urethane polymer to wrap around the entire body.

### Parylene Coating of the OLED Device

Dichloro (2,2) paracyclophane monomer (Dix-C, PARYLENE JAPAN, Tokyo, Japan) was vaporized by heating to 150°C under reduced pressure to 3.0 Pa using a chemical vapour deposition apparatus, a system of interconnected components (monomer vaporisation chamber - heating furnace - deposition chamber) fabricated by our laboratory (K.F.). The vaporized monomer was then transferred to a chamber set at 680°C and decomposed into monomeric diradicals. The monomers were reheated to 150°C and vaporized again. This vapour was then diffused into a room-temperature chamber in which the OLED devices were placed. The monomers were polymerized, resulting in a thin film of poly (p-xylylene) derivative (Parylene C). After a 1 μm-thick Parylene C film was accumulated, the OLED devices were rotated and coated with an additional 1 μm-thick Parylene C film to minimise coating non-uniformity. By this method, the edges of the devices were also adequately coated.

### Fabrication of the Wireless Power Transmission Platform

The newly developed wireless power transmission platform consists of a high-frequency power generator, a power amplifier (PT200A, Pleiades Technologies, Fukuoka, Japan), a power meter (SX200, DAI-ICHI DENPA KOGYO, Tokyo, Japan), and a container holding an antenna transmitting a 13.56 MHz radio wave (PTA-A3, Pleiades Technologies, Fukuoka, Japan). The container was box-shaped and its dimensions were configured to be 340 (W)×500 (L)×160 (H) mm (internal dimensions) and 440 × 600 × 170 mm (external dimensions), being sufficient to house a standard cage (bottom dimensions: 206 (W) × 365 (L) mm, uppermost dimensions: 263 (W)× 426 (L) mm, height: 202 mm) for multiple rats.

### Orthotopic Tumor Model

Six-week-old male Sprague Dawley (SD) rats (186 - 202 g) (Japan SLC Inc., Japan) residing within the SPF area were used. To establish an orthotopic hepatoma model, SD rats were administered a compound anesthetic [medetomidine (0.3 mg/kg) (Nippon Zenyaku Kogyo, Japan), midazolam (4.0 mg/kg) (Sandz Corp., Japan), and butorphanol (5.0 mg/kg) (Meiji Seika Pharma, Japan)]. A minor laparotomy was performed, allowing the left lobe of the liver to be exposed, followed by an injection of 30 μL of N1-S1 (A rat hepatoma cell line, ATCC, CRL-1604, Manassas, VA) cell suspension (2.0×10^5^ cells) in phosphate-buffered saline into the subhepatic capsule with a 30 G needle. Tumor size was gauged using an ultrasonic imaging system (10 MHz probe, VEVO770, VISUAL SONIC, Toronto, Ontario, Canada). Individual rats exhibiting tumor volumes between 20 and 30 mm³, as determined on the fourth day post-implantation, were selected for inclusion in the experiments. All procedures were performed in compliance with the Science Council of Japan’s Guidelines for the Proper Conduct of Animal Experiments, with the experimental protocols receiving approval from the National Defense Medical College Animal Care and Use Committee (Approval No. 19009, 23045).

### Implantation OLEDs in Animals

Subsequent to the intraperitoneal administration of the compound anesthetic, laparotomy was performed to expose the left lobe of the liver. The location of the tumor was verified via an ultrasound system, and the OLED device was positioned so that the centre of the tumor was aligned with the centre of the OLED’s emitting surface. Medical-grade cyanoacrylate adhesive (5 μl) (Aron alpha A, Daiichi Sankyo, Japan) was applied to the OLED’s four corners to secure the device affixed onto the liver.

### Photosensitizer (PS)

Temoporfin (5,10,15,20-tetrakis (3-hydroxyphenyl) chlorin; mTHPC) (SML1707, Sigma-Aldrich) was dissolved in anhydrous ethanol (443611, Sigma-Aldrich) and then in propylene glycol (3980039, Sigma-Aldrich) to make a 1 mg/ml solution (anhydrous ethanol : propylene glycol = 9 : 14). The solution was diluted 5-fold with Milli-Q water and administered through the tail vein of each rat (0.5 mg/kg).

For the measurement of absorbance spectra of the PS, a 0.2-mg/ml aqueous solution of mTHPC was prepared, and the absorbance was measured using a spectrophotometer (JASCOV630; Shimadzu Corporation, Tokyo, Japan).

### Metronomic PDT in the Orthotopic Tumor Model

After the tumor reached a certain size (20 - 30 mm^3^) (approximately 4 days after inoculation of cancer cells), mTHPC was administered via the tail vein (0.5 mg/kg) and an OLED device was affixed onto the liver so that the OLED’s emitting surface covered the tumor surface. The rats were then transferred to a cage in the antenna board. Three hours after administration of the PS, light emission was started and continued for 48 hours. The transmitter power was set at 8 W for both green and red OLEDs. For the group with administration of the PS twice, an additional dose of mTHPC (0.5 mg/kg) was administered via the tail vein and light emission was continued for another 48 h. After completion of mPDT, the tumor size was measured by an ultrasound imaging system and then the rats were euthanised and the left liver lobe of each rat was extracted.

## Acknowledgments

We thank K. Aoki and Y. Mitsui for their technical assistance; Dr. T. Fujie (Tokyo Institute of Technology, Japan) and Dr. K. Yamagishi (SUTD, Singapore) for discussions; and the staff of the R&D Dept. of MORESCO Corporation (Kobe Japan) for providing sealing resin. Y. Mo. is supported by JSPS KAKENHI Grant (21H03846, 21K19935), Fukuda Foundation for Medical Technology, The Uehara Memorial Foundation, Suzuken Memorial Foundation, and Takeda Science Foundation. K. F. is supported by JST A-Step Grant (JPMJTM20GY), Kyutec Research Grant, and Tateishi Science and Technology Foundation (B2221904). K. M. is supported by JSPS KAKENHI Grant (19H02599).

## Author Contributions

Conceptualization and experimental design: Y.I., K.S., K.F. and Y.Mo. Methodology: Y. I., K.S., K.F. and Y.Mo. Investigation: Y.I., I.K., T.T. and T.S. (animal experiments), Y.I. and T.T. (ultrasound imaging), Y.I. (immunocytochemistry and imaging), K.S., K.F., K.E., Y.Mi. and R.Y. (fabrication of the OLED device and wireless power transmission system), K.K., N.K., T.G. and K.M. (assembly of OLED elements), K.F. (parylene coating), Y.I. and Y.Mi. (measurement of optical properties), Y.I. (data analysis). Funding acquisition: K.F., K.M. and Y.Mo. Supervision: K.S., K.F., H.T. and Y.Mo. Writing-original draft: Y.I. and Y.Mo. Writing-review and editing: Y.I., K.S., K.F. and Y.Mo.

## Additional Methods

### Measurement of Optical Properties of the OLED Device

The wavelength spectrum of the light emitted from the OLEDs (Figure 2f) was measured using a photonic multichannel analyser (PMA-12, HAMAMATSU PHOTONICS K.K. Hamamatsu, Japan). For luminance analysis, a matrix of 45 measurement nodes was constructed at 1-mm intervals on the luminous plane (Figure S1a), and the luminance at each node was measured using a luminance meter (LS-160, KONICA MINOLTA, Tokyo, Japan) at an aperture size of 0.5 mm φ. The angular luminance of the emitting surface (angle-dependent light distribution characteristics) (Figure S7) was measured using the luminance meter with an aperture size of 0.2 mm φ. For luminance distribution analysis (Figure 2d), a luminance image was captured by a camera integrated with a CMOS sensor (EOS Kiss ×10, Canon, Tokyo, Japan) and visualized using Python 3.10, preprocessing with pandas 1.4, computing with numpy 1.23, and creating graphs with matplotlib 3.5.

### Measurement of the Spatial Distribution of Electromagnetic Field Strength

The homogeneity in the distribution of the electromagnetic field strength of the power transmission platform was assessed in the following way. The OLED device was held above the antenna board and the emitted light intensity (irradiance) was measured at different heights from the base of the antenna board. Specifically, the OLED device was placed at five different heights, 35 mm, 60 mm, 85 mm, 110 mm and 135 mm, from the antenna board, and the horizontal distribution of light intensity was examined at each height (Figure 3c). The transmitter power was set to 8 W. A power meter with a photodiode sensor (PD300-BB, Ophir, Saitama, Saitama, Japan) was used to measure the light intensity.

On the other hand, for animal experiments, only OLED devices with an irradiance within a certain range (400 - 450 μW/cm^2^) were used, measured at the center position of the antenna board and 60 mm from the bottom of the antenna board.

### Measurement of Heat Generated by the OLED Device

In order to assess the unavoidable thermal generation from the operation of the implanted OLED device on the liver surface *in vivo*, the device was adhered on the left lobe, and the temperatures of the device surface and the liver surface were measured by infrared thermography (FSV-2000, Apiste, Osaka, Japan). The heat from the device operation was mainly due to the power-receiving coil.

### Estimation of the Light Energy Received by One Animal by mPDT

A study was conducted to estimate the total amount of light energy received by one animal during mPDT. A spacer was placed between the cage floor and the antenna board so that the height level of the rat’s liver in the prone position was 60 mm from the bottom of the antenna board. To trace movement of a rat, an LED chip driven by near-field communication (KP-NFLEG (λ=530 nm), Kyoritsu Electronics Industry, Osaka, Japan) was implanted subcutaneously in the posterior neck of the rat and the emission from the LED was filmed by a video camera (EOS Kiss × 10, Canon, Tokyo, Japan) placed above the cage (Figure S4a). The trajectory of the rat’s movement was analysed using UMA tracker (Yamanaka and Takeuchi, 2018), an open-source software for behavioural analysis. Total light energy received was calculated by multiplying the time spent by the rat in each area of the cage floor by the irradiance of the OLED device at that location (Figure S4b, c, d, e).

### Cell Line

A rat hepatoma cell line (N1S1, ATCC, CRL-1604, Manassas, VA, USA) was employed for the experimental procedures. Cells were propagated in 10% FBS, penicillin (100 U/mL) (Thermo Fisher Scientific, Japan), streptomycin (100 μg/mL) (Thermo Fisher Scientific, Japan), and amphotericin B (0.25 μg/mL) (Sigma-Aldrich K.K., Japan) within Dulbecco’s Modified Eagle’s Medium and cultured in a temperature-controlled incubator at 37°C with 5% CO_2_ and 95% air.

### Evaluation of Temporal Accumulation of the PS in the Tumor

The PS (mTHPC, 0.5 mg/kg) was administered intravenously 7 days after inoculation of cancer cells, and the left liver lobes including the tumors were extracted at 12, 24, 48, and 72 hours after PS administration (n=1, respectively). The tissue samples underwent perfusion with saline and 4% paraformaldehyde, followed by freezing at −80°C. Thin sections of 5 μm in thickness were then prepared and observed using a fluorescence microscope (BZ-X 700, KEYENCE, Japan). Autofluorescence was detected at Ex. 470 nm / Fluo. 500-550 nm and mTHPC was detected at Ex. 405 nm / Fluo. 620-660 nm. Exposure times were all 0.5 s.

To investigate the concentration of mTHPC in tissues, the intensity of mTHPC-fluorescence in the liver and tumor tissue was measured and the intensity per unit area was calculated using ImageJ software (https://imagej.net/ij/index.html).

### Tumor Volume Measurement and Histopathological Examination

Tumor dimensions were determined by using an ultrasonic imaging system by the following formula: long diameter × short diameter × height × π/6.

For hematoxylin and eosin (H&E) staining and terminal deoxynucleotidyl transferase (TUNEL) staining, the specimens were prepared according to the respective staining procedures after 10% formalin fixation and paraffin-embedded sections were prepared. Upon sectioning and staining, images were digitally captured at 400× resolution utilizing a digital scanner (Nano Zoomer 2.0 HT; Hamamatsu Photonics, Shizuoka, Japan). For image analysis, QuPath Bioimage analysis v.0.2.3, an open-source software designed for digital pathology and whole slide image analysis (Irish Molecular Pathology Laboratory, Centre for Cancer Research and Cell Biology, Queen’s University Belfast), was used.

Apoptotic fractions were calculated as the ratio of the manually marked apoptotic area to the total tumor expanse within the tissue section that had the highest percentage of tumor in serial sections. These apoptotic fractions were then normalized by dividing by the total tumor volume (normalized apoptotic fractions[1]).

The density of dead cell nuclei per area at every 0.25-mm depth from the tumor surface was calculated from high magnification images (Figure 5i).

### Evaluation of Adverse Events

Rats were weighed before and after mPDT. To evaluate hepatic function, blood was collected from the tail vein before and after mPDT, and the hepatic excretory enzymes alanine aminotransferase (ALT) and aspartate aminotransferase (AST) were quantified.

### Statistical Analysis

Data are presented as means ± standard deviation. Statistical analysis was conducted using GraphPad Prism 9 (GraphPad Software, California, USA). Tumor volume was analysed via one-way analysis of variance with between-group comparisons performed using Tukey-Kramer’s HSD test. Thickness of the capsule, width and depth of the apoptotic area and normalized apoptotic fraction, AST, ALT, and body weight were analysed via a nonparametric Mann-Whitney U test. The density of dead cell nuclei was analysed via two-way analysis of variance, with between-group comparisons performed using Šídák’s multiple comparisons test. A p-value less than 0.05 was considered statistically significant.

---

**Supplementary Table 1.** Effect of ferrite sheets on Q-factor in the OLED device. The use of OLED elements increased the resistance (R) of the power-receiving circuit and decreased the Q-factor. However, the use of ferrite sheets prevented the Q-factor from decreasing even in the presence of OLED elements, thus preventing a decrease in transmission efficiency.

|  | OLED (-) |  | OLED (+) |  |
| --- | --- | --- | --- | --- |
|  | ferrite (-) | ferrite (+) | ferrite (-) | ferrite (+) |
| inductance ($\mu\text{H}$) | 0.84 | 1.37 | 0.84 | 1.37 |
| R ($\Omega$) | 1.53 | 2.81 | 1.53 | 3.06 |
| Q-factor | 46.8 | 41.5 | 10.3 | 38.2 |

**Figure S1.**
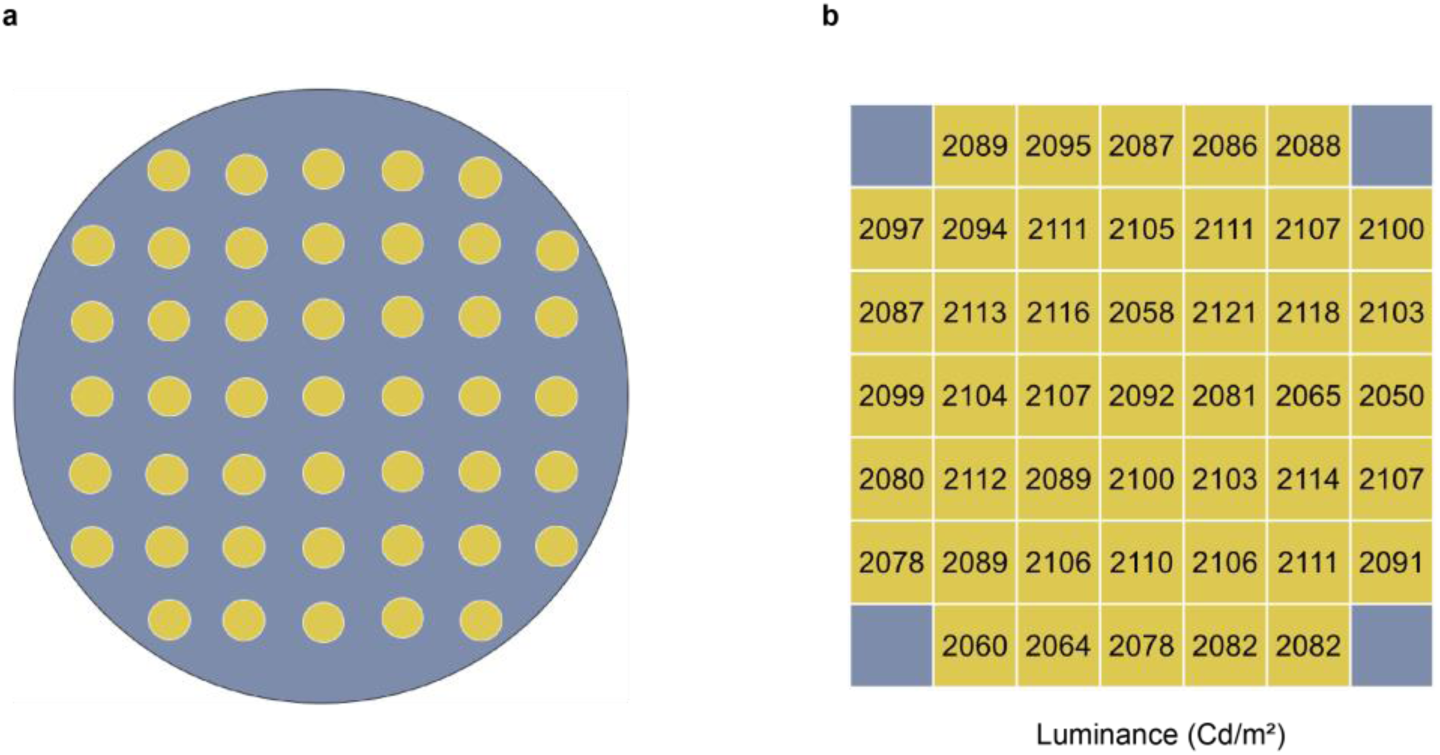
Homogeneously emitting light from the OLED. a) Visual representation of the OLED emitting surface and positions of the measurement nodes. b) Irradiance values at the nodes. The variation is within ± 0.8%, indicating highly homogeneous irradiance characteristics. The OLED device (green) was operated at a transmitter power setting of 0.6 W. a.u, arbitrary units.

**Figure S2.**
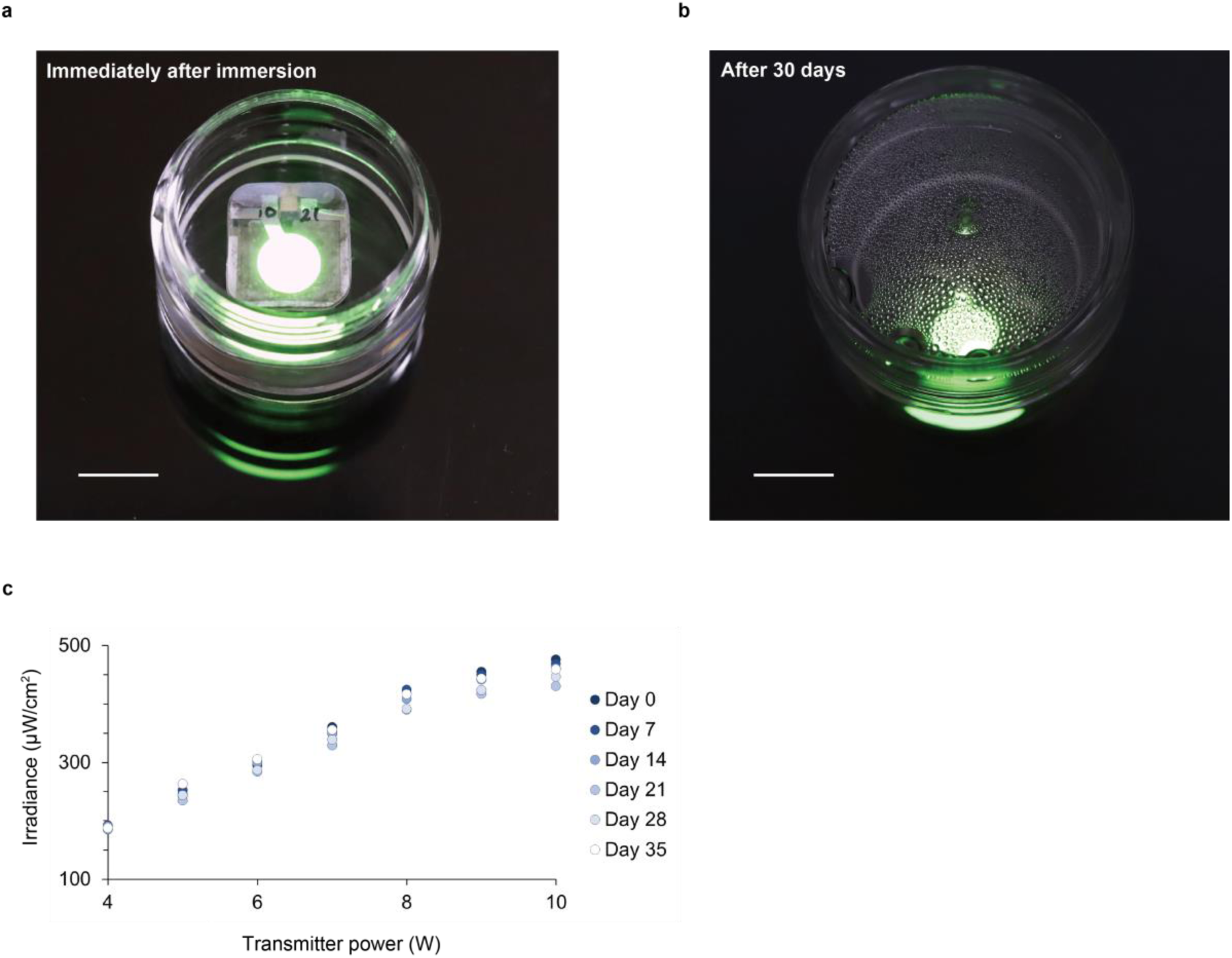
High degree of water sealing by Parylene C coating. OLED (green) emitting light in phosphate-buffered saline: immediately after immersion (a) and 30 days after immersion (b). c) Device irradiance variation over time. Encasing the OLED (green) in Parylene C allowed it to emit light continuously in phosphate-buffered saline for over one month, thereby preserving its water sealing capabilities. Scale bar = 10 mm.

**Figure S3.**
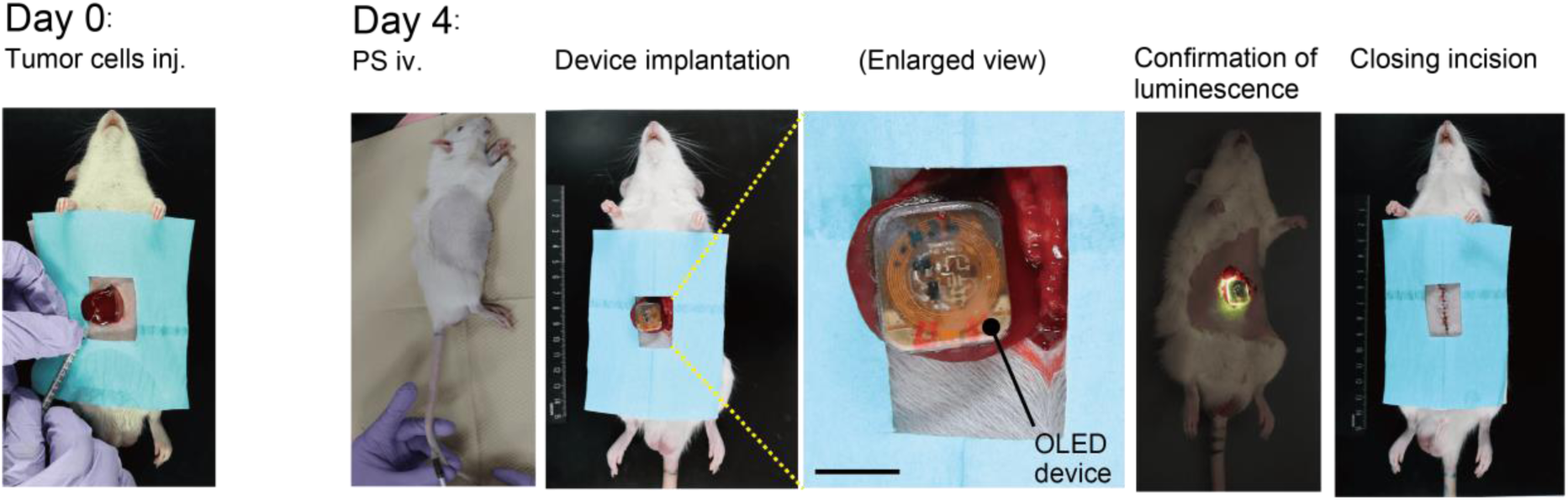
Orthotopic rat hepatoma model and OLED device implantation. Sequential steps illustrating the construction of a hepatoma model in a rat. Day 0: Injection of tumor cell suspension into the hepatic lobe (Tumor cells inj.). Day 4: Intravenous administration of the photosensitizer via the tail vein (PS iv). This is followed by the surgical implantation of an OLED device (Device implantation) and confirmation of the device’s luminescence (Confirmation of luminescence). The liver is returned to its original position with the OLED attached and then the skin wound is closed (Closing incision).

**Figure S4.**
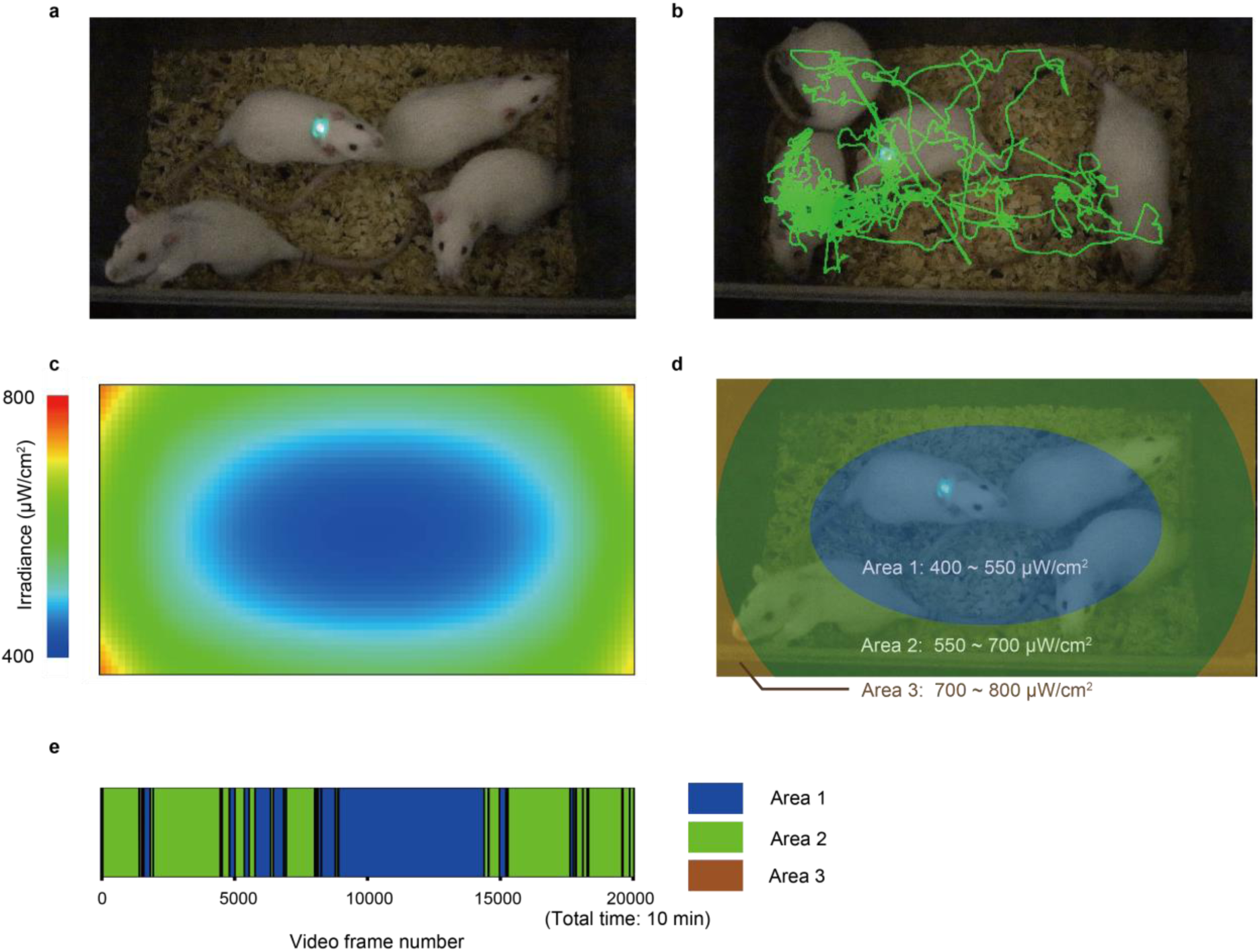
Estimation of the light energy received by one animal by mPDT. a) LEDs driven by near-field communication (NFC) were implanted subcutaneously in the posterior necks of rats to serve as markers for movement tracing. b) The trajectory of one rat’s 10-minute movement. c) Distribution of OLED irradiance at a height of 60 mm from the bottom of the antenna board (simplified while simulating the irradiance distribution in Figure 3c). d) To facilitate the calculation of estimated energy received, the floor surface of the animal cage was divided into three sections based on the irradiance distribution of (c). e) Behavioural mapping of a rat over a 10-minute period (20000 video frames), showing where the individual stayed in the three divided areas on the cage floor. In this case, the percentages of time spent in the areas were 38% in Area 1, 62% in Area 2 and 0% in Area 3. Therefore, the irradiance per unit time (sec) for this 10-minute period can be calculated to be 493-643 μW/cm^2^. On this basis, and assuming a stay of 48 hours, the total energy received by the individual can be estimated to be between 85 and 111 J/cm^2^. A green OLED device was used and the transmitter power setting was 8 W.

**Figure S5.**
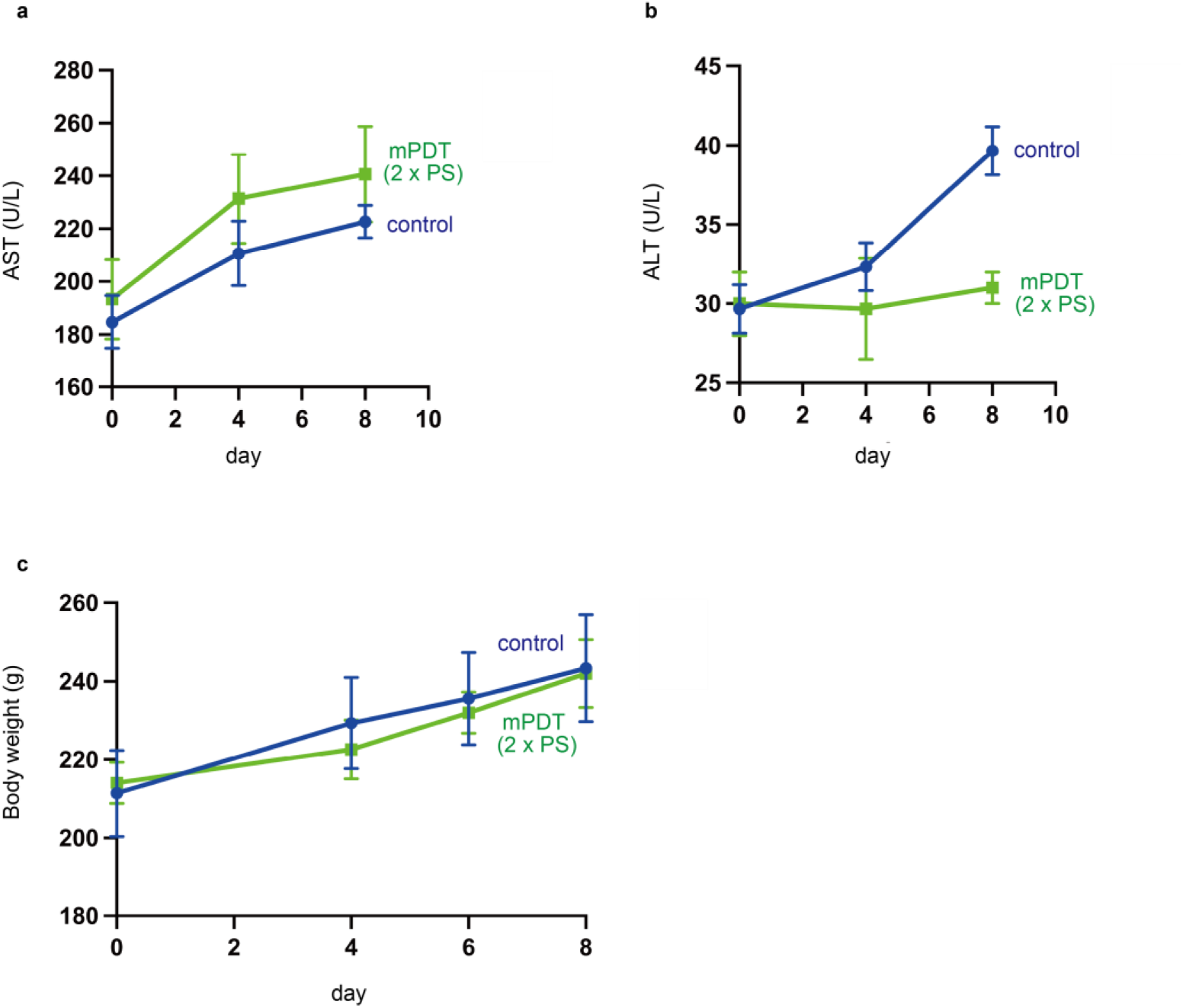
No adverse effects from OLED-based mPDT. Changes in aspartate aminotransferase (AST) (a) and alanine aminotransferase (ALT) levels (b) during mPDT using green OLED devices (n=3, each) (median AST on day 8: mPDT: 248 U/L, control: 224 U/L, p=0.4; median ALT on day 8: mPDT: 31 U/L, control: 40 U/L, p=0.1). c) Changes in body weight during mPDT (n=5, each) (median weight on day 8: mPDT: 246 g, control: 252 g, p=0.397).

**Figure S6.**
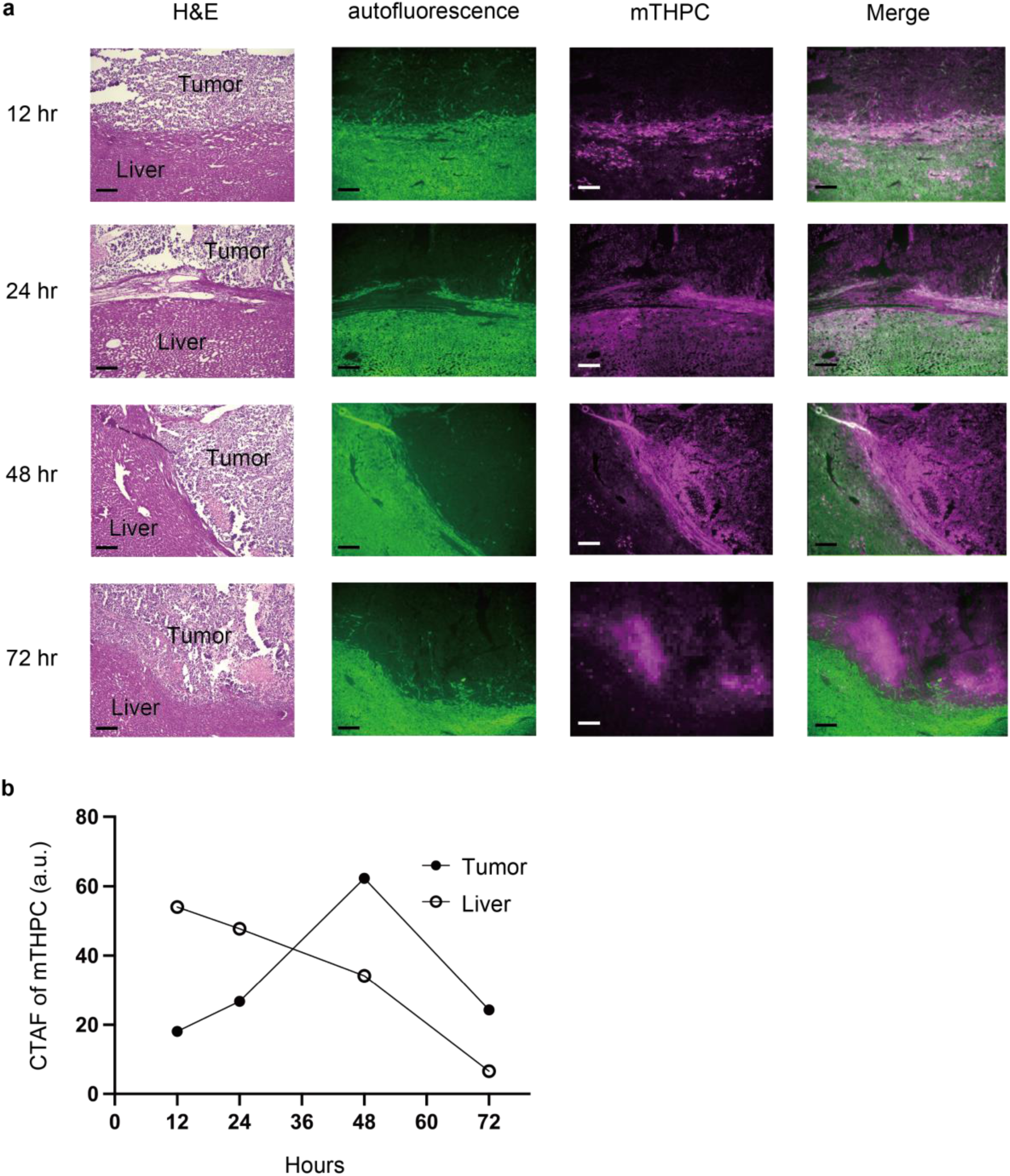
Changes in PS accumulation in the tumor over time. a) Tissues extracted at 12, 24, 48 and 72 hours after PS administration. b) Changes in PS concentration in tissues over time. The mTHPC concentration in the tumor increased over time, peaking at 48 hours after administration, at which time it exceeded the mTHPC concentration in the surrounding normal liver. However, at 72 hours after administration, the mTHPC concentration in the tumor was decreased and was almost the same as the level at 12 hours after administration. The tissue concentration of mTHPC was substituted by corrected total area fluorescence (CTAF), which was calculated using the following formula: CTAF = [Total fluorescence in the ROI - Total fluorescence in the background region of exactly the same area as the ROI]/area of the ROI. ROI is an abbreviation for region of interest. a.u, arbitrary units. Scale bar = 0.2 mm.

**Figure S7.**
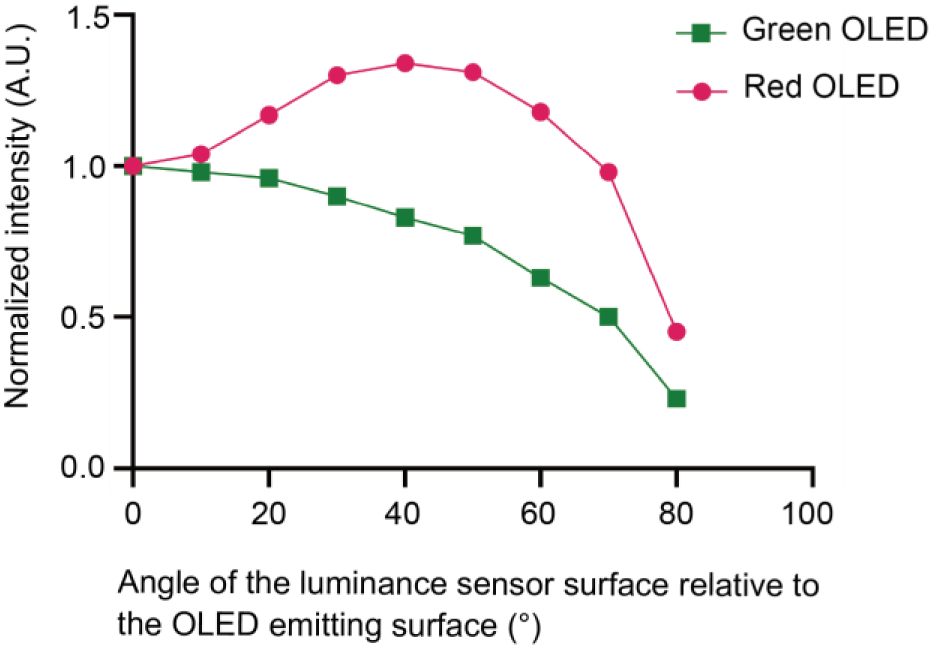
Angle-dependent light distribution characteristics of OLED elements. Measurements were taken at different angles of the luminance meter receiving surface relative to the OLED emitting surface. Horizontal (0°), Vertical (90°).

**Legend for Supplemental Movie 1 |**

An OLED attached to the liver emitting light under wireless drive.

**Legend for Supplemental Movie 2 |**

An OLED attached to the liver wirelessly emitting light in the abdominal cavity, visible through laparoscopy. The OLED’s illuminating surface was positioned upside down for easy observation of its light.

